# Local climate and habitat continuity interact to alter contemporary dispersal potential

**DOI:** 10.1101/2020.04.17.047050

**Authors:** Lauren L. Sullivan, Zoe M. Portlas, Jill A. Hamilton

## Abstract

Understanding the evolution of dispersal under changing global environments is essential to predicting a species ability to track shifting ecological niches. Two important, but anthropogenically altered, sources of selection on dispersal are climate and habitat continuity. Despite the likelihood these global drivers of selection act simultaneously on plant populations, their combined effects on dispersal are rarely examined. To understand the interactive effect of climate and habitat continuity on dispersal potential, we study Geum triflorum - a perennial grassland species that spans a wide range of environments, including both continuous prairie and isolated alvar habitats. We explore how the local climate of the growing season and habitat continuity (continuous vs isolated) interact to alter dispersal potential. We find a consistent interactive effect of local climate and habitat continuity on dispersal potential. Across continuous prairie populations, an increased number of growing degree days favors traits that increase dispersal potential. However, for isolated alvar populations, dispersal potential tends to decrease as the number of growing degree days increase. Our findings suggest that under continued warming, populations in continuous habitats will benefit from increased gene flow, while isolated populations will become increasingly segregated, with reduced potential to track shifting fitness optima.

## Introduction

Altered global environments promote the evolution of dispersal as a means for species to track shifting fitness optima (Kokko and López-Sepulcre 2006; Aitken et al. 2008). Indeed, dispersal evolves even in the simplest of systems in response to altered conditions (Hamilton and May 1977; Hargreaves and Eckert 2014), and can be especially important for sessile organisms, such as plants, that have limited opportunities for movement. Seed dispersal, or the movement of offspring away from the source parent plant, provides the opportunity for migration (Zobel et al. 2010), facilitating gene flow (Sexton et al. 2009), range shifts (Davis and Shaw 2001; Hargreaves et al. 2015), and spatial tracking of favorable environmental conditions (Edelaar and Bolnick 2012; Brehm et al. 2019) - all which allow for species persistence despite local extirpation (Tilman 1994). Two important drivers of selection on dispersal ability are climate and habitat connectivity or isolation (Mantyka-Pringle et al. 2012). Despite the likelihood that these factors synergistically impact plant populations, they have rarely been studied empirically in concert (but see: Delattre et al. (2013)). However, as both climate change and alterations to land-use impact the spatial arrangement and quality of available habitat (Eriksson and Jakobsson 1999; Rockström 2009), it is imperative to determine how these factors interact to influence the evolution of dispersal and potential connectivity across increasingly heterogeneous landscapes.

Variation in the local thermal environment can have both positive and negative effects on dispersal capacity, altering plant species’ ability to track changing climatic conditions (Corlett and Westcott 2013). Warming temperatures have been associated with intraspecific increases in plant height (Bjorkman et al. 2018), directly increasing potential dispersal distance (Thomson et al. 2011; Teller et al. 2014). However, for some species, ranges are contracting as climates warm, indicating that increased temperatures do not necessarily impact species dispersal potential in the same manner (Zhu et al. 2012). Adaptation to local conditions can also play a role in the evolution of dispersal potential. Hargreaves et al. (2015) used simulations to demonstrate the relationship between adaptation to local environments and dispersal ability. Where genotypes were adapted to locally stable climate optima, they exhibited reduced dispersal ability, selecting against dispersal in order to remain in environments associated with increased relative fitness (Hargreaves et al. 2015). This selection against dispersal could limit species’ ability to track shifting optima, particularly if the trait variation needed for longer-distance movement does not exist. Thus, understanding how local climates can alter species dispersal trait variation will impact our ability to predict species’ response to changing conditions, particularly the potential for range shifts and the maintenance of connectivity in a warming climate (Parmesan and Yohe 2003; Parmesan 2006).

Multiple predictions also arise for habitat continuity as a selective force influencing dispersal trait variation. Classic theory from island biogeography predicts that isolated island populations will exhibit reduced dispersal relative to mainland populations in order to limit dispersal beyond suitable habitats (MacArthur and Wilson 1967). Habitat fragmentation can have similar effects on selection for dispersal ability (Fahrig 2003, 2017). Experimental studies have shown that habitat fragmentation can promote rapid evolution of both long-distance (Williams et al. 2016), and shorter-distance (Cheptou et al. 2008) dispersal. These differences likely depend on the scale of fragmentation, as the distance to suitable habitat may select for varying dispersal strategies (Cody and Overton 1996; Baguette et al. 2012). Travis & Dytham (1998) explored these mechanisms with simulation studies and found that as habitat availability was reduced, selection favored a reduction in dispersal, particularly as the likelihood of reaching suitable habitat decreased. However, they also found that as habitat autocorrelation increased and the likelihood similar habitat patches being adjacent increased, selection favored longer-distance dispersal. Therefore, the degree of fragmentation – which includes both the spatial arrangement and amount of suitable habitat – will impact selection on trait variation associated with dispersal ability, and consequently the evolution of dispersal across heterogeneous environments.

Here, we examine the relationship between local climate and habitat continuity to test their impact on dispersal evolution using a species that spans broad climatic and isolation gradients, *Geum triflorum*, Prairie Smoke (Rohrer 1993). We examine contemporary differences in population-level dispersal trait values to test for alterations in the trajectory of dispersal evolution. Wind-dispersed plant species, like *G. triflorum*, have evolved multiple mechanisms that promote movement by wind, including structures such as plumes, wings, or samaras that increase time aloft and in turn increase dispersal distances (Greene and Johnson 1990; Lentink et al. 2009; Varshney et al. 2012). Population-specific variation in dispersal traits would indicate that selection likely contributes to regional differences, influencing the evolutionary trajectory of dispersal traits across the range of *G. triflorum*. Specifically, we aim to determine if alterations to local climate (specifically, the number of growing degree days above 5°C), and habitat continuity (either continuous or isolated regional habitats) have positive, negative or interactive effects on contemporary dispersal trait variation including shape, mass, and falling speed that collectively influence dispersal potential. We measured dispersal traits from 100 *G. triflorum* populations across its range, including populations from both historically continuous and isolated habitat patches. Using these traits, we parameterized dispersal models to determine dispersal potential of individuals within each population. We found that local climate and habitat continuity interact to influence dispersal trait variation, and in turn dispersal potential of *G. triflorum*. In continuous habitats, increasing the number of growing degree days increased dispersal ability; while in isolated habitats, increasing the number of growing degree days reduced dispersal ability. These data indicate the importance of considering the interaction between local climate and spatial structure of populations in predicting species’ ability to spatially track shifting fitness optima under global change.

## Materials and Methods

### Study Species

*Geum triflorum* (Pursh), or Prairie Smoke (Rosaceae) is an herbaceous perennial distributed across much of northern North America (US and Canada) and in the southwestern United States and California (Gajewski 1958; Rohrer 1993). Inflorescences typically have three nodding flowers that become erect at seed set. The sessile infructescence contains multiple achenes, each connected to a densely wooly style that remains intact during dispersal (Fig. 1a). This structure, including both the achenes and style (Fig. 1b-d), is termed the diaspore. These styles likely promote wind dispersal by increasing the time-aloft for diaspores following release from the maternal plant (Greene and Johnson 1989, 1990).

**Figure 1:**
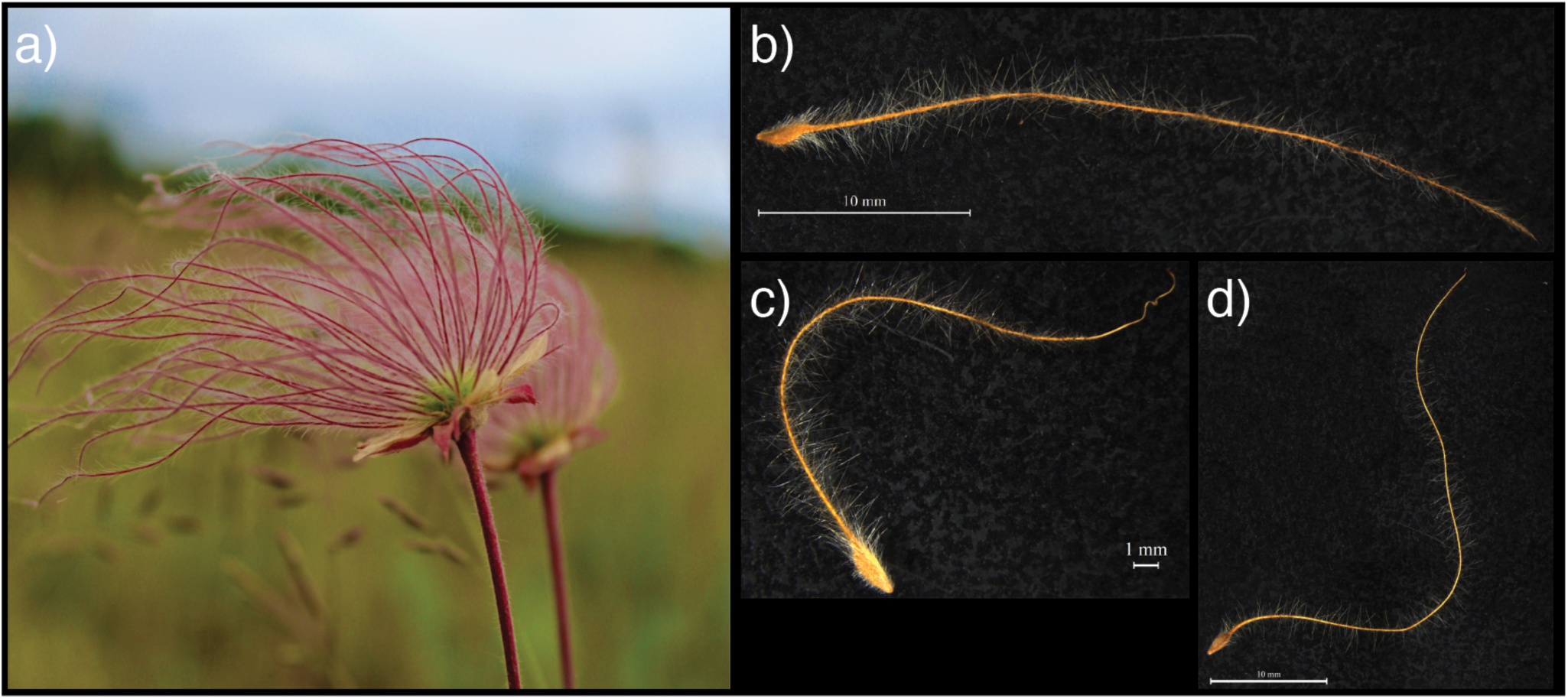
*Geum triflorum* A) fruiting head and B-D) diaspores. Each diaspore contains an achene that is connected to a densely wooly style that remains intact at the time of dispersal. These hairs help slow the rate of falling, and thus help increase the dispersal ability of these diaspores. Panels B-D represent the range of diaspore shapes and sizes.

### Sampling

We collected individual seed heads from populations of *G. triflorum* in 2002, 2003, 2015, and 2016. These populations were defined as distinct groups of plants separated from other populations by at least 1km. Individual seed heads were collected from between 40 to 150 plants along a 100m transect in 2002, 2003, and 2015 and along a 1000m transect in 2016. See Hamilton and Eckert (2007) for detailed sampling methods. In total, we sampled 60 continuous prairie populations and 40 isolated alvar populations (ecoregional types described in ‘*Habitat Continuity’* section below), all of which exhibit similar ranges of climatic variation.

### Habitat Continuity

To determine how habitat continuity altered dispersal traits, we sampled *G. triflorum* populations in both its historically continuous range throughout the midwestern prairies of North America, and across disjunct populations geographically isolated on limestone barrens known as alvar habitats, which are distributed throughout the Great Lakes region and into Manitoba, Canada. The species occurs in four of seven alvar regions across the Great Lakes (Catling and Brownell 1995; Reschke et al. 1999); the Napanee Plain, the Carden Plain, Western New York, and Manitoulin Island, as well as isolated alvar habitats on Drummond Island, Michigan and northern Manitoba. *G. triflorum’s* prairie range extends from west of the Great Lakes across the Rocky Mountains, covering much of the Great Plains of Canada and the United States.

Genetic connectivity varies between prairie and alvar populations (Hamilton and Eckert 2007), which has implications for the evolution of dispersal traits. Prairie populations have historically experienced higher levels of genetic connectivity (Hamilton and Eckert 2007), but contemporary fragmentation over the last century (e.g. Wright and Wimberly 2013; Lark et al. 2018) has led to increased isolation. In contrast, historically isolated alvar populations are not only disjunct from the main contiguous range across the midwestern prairies, but also are isolated from each other. Genetic data indicate that alvar populations have a subset of the genetic variation found in prairie populations, and within the same geographic distance alvar population pairs are significantly more genetically differentiated from each other than prairie population pairs (Hamilton and Eckert 2007). Thus, contemporary differentiation between prairie and alvar populations likely reflects a combination of colonization history and contemporary gene flow, stochastic processes associated with shifts in population demography, and changing selective processes leading to adaptation across regional environments.

### Climate

To determine how climate influenced dispersal traits of *G. triflorum*, we used year of collection, latitude, longitude, and elevation from sampling locations as inputs into ClimateNA (Wang et al. 2016) to estimate annual climate variables based on geographic coordinates. For each population, we were able to estimate annual climate variables including many related to precipitation and temperature. In order to reduce redundancy and account for correlation across climate variables we performed a principle component analysis (PCA). We found temperature variables (number of growing degree days above 5oC, mean annual temperature, frost free period, etc) largely loaded on the PC1 axis and explained 51% of the variation, while variables associated with water-availability (mean annual precipitation, climatic moisture deficit, annual heat moisture index, etc.) largely loaded on PC2 axis and explained 23.6% of the variation (Appendix S1). To simplify our analyses, we described climate using the “number of growing degree days above 5oC” variable (hereafter referred to as DD5) as it loads highly on PC1 representing much of the temperature variability spanning our populations. Furthermore, it is a biologically meaningful climate variable as it reflects the required heat sum associated with the onset of growth (Beaubien and Hamann 2011). It is predicted to be particularly sensitive to climate change (McGinn and Shepherd 2003) as shifts in DD5 for spring perennials, such as *Geum triflorum*, may have substantial influence on the timing and duration of the growing season (Beaubien and Hamann 2011; Whittet et al. 2017). Note: we also ran our analyses with the precipitation variables that loaded highly on PC2, however the results were rarely significant.

### Dispersal Trait Measurements

To measure dispersal traits, we randomly selected three individuals, each representing one maternal family, from a population. We then measured three types of dispersal traits on all diaspores (Fig. 1), including mass, morphology, and terminal velocity (or falling speed). Pooling five individual diaspores per maternal family, we first estimated total diaspore mass. We weighed both the achene and style separately to the nearest 0.0001g. Following this, we examined morphological variation for each of five individual diaspores per maternal family. We photographed diaspores on a 5mm2 grid using a Leica DM2500 dissecting microscope for all morphological measurements. We placed a glass sheet atop each diaspore to uniformly flatten the dispersal structures. All measurements were made using ImageJ (Schneider et al. 2012). We measured the total length of the style and achene to the nearest 0.001mm, and the achene area to the nearest 0.001mm2. We measured the total length of the style and achene, which included the entire path length, not “folded” length, and achene area was calculated using the length and width measured at the longest and widest points, respectively. We also calculated diaspore shape index, which is a measure of 2D shape of each diaspore. Finally, to measure terminal velocity, we dropped up to five individual intact diaspores per maternal family through a modified 10cm wide PVC tube with two sets of light sensor arrays spaced 57cm apart that record when the diaspores passed through both sets of arrays. We recorded the time it took the diaspores to cross this distance as the terminal velocity. For a full description of the terminal velocity measurement device and setup, see Sullivan et al. (2018). Diaspores were dropped from a height of 1m to ensure they reached terminal velocity. For terminal velocity measurements we used data from 26 prairie populations and 17 alvar populations reflecting contemporary collections from 2015 and 2016.

In order to translate terminal velocity measurements into dispersal ability, we used the WALD model (Katul et al. 2005), which has been empirically validated for grassland species (Soons et al. 2004; Sullivan et al. 2018), to calculate the dispersal distances traveled by the farthest 1% of dispersers. To do this, we followed Sullivan et al. (2018) and used plant traits including height at diaspore release and terminal velocity to parameterize the estimated dispersal kernel for each diaspore. We estimated pooled height at diaspore release for each region as the average height from 53 prairie and 18 alvar flowering stems of *G. triflorum* (measured from rosette to base of sepals) from herbaria specimens provided by the University of Minnesota (MIN), University of Manitoba (WIN), the Canadian Museum of Nature (CAN), and the Agriculture and Agri-Food Canada National Collection of Vascular Plants (DAO). We estimated the canopy height to be 0.2m for both regions at the time of *G. triflorum* seeding, as this is early in the season and many plants are still short. Finally, we parameterized wind values from weather station readings near Moorhead, MN. We used a grand average of 7-day wind averages collected approximately every 4 days during the month of June in 2017 and 2018 as this is when the majority of *G. triflorum* diaspore dispersal occurs. We used these data to estimate wind parameters in prairie systems, and assumed the wind to be 50% of this value for alvars, as these isolated habitats are surrounded by forest trees, which can substantially reduce wind effects (Damschen et al. 2014). Dispersal kernels were modeled for each individual diaspore within each region, following which we extracted the predicted distance travelled if the diaspore achieved its long-distance dispersal potential (moved to the farthest 1% of its dispersal kernel - Higgins et al. 2008).

### Statistical Analysis

To determine how habitat continuity (alvar vs prairie) and local climate interact to influence dispersal traits of *G triflorum*, we used R v3.5.0 (R Core Team 2018) to conduct mixed-effects models with the lmer() function in the lme4 package (Bates et al. 2015), and used the lmerTest package (Kuznetsova et al. 2014) to extract p-values. We also used the MuMIn package (Barton 2018) to determine the contributions of the fixed and random effects using r2 values. Dependent variables examined were the three categories of dispersal traits, including mass, morphology, and terminal velocity. All trait variables were transformed using a square root transformation to meet the assumption of normality. Additionally, to determine the effect of local climate and habitat continuity on dispersal potential, we analyzed the log of the distance travelled by the farthest 1% of dispersing individuals. In all models, the fixed effects included the interaction between geographic region (prairie vs alvar) and the log DD5 estimated from the growing season in which diaspores were collected. Random effects included replicates within families when appropriate, as well as year to account for differences in years of trait collections. Code and data are available in the supplemental information (Appendix S2, S3).

## Results

### Mass Measurements

We found a significant interaction between habitat continuity (alvar vs prairie) and local climate for total diaspore mass (Table 1a, *p*=0.021), where fixed effects explained 4.6% of the variance, and fixed and random effects together explained 65.6% of the variance. As the length of the growing season increases, prairie diaspores become heavier, while alvar diaspores become lighter (Fig. 2a).

**Table 1.** Mixed effects model results for A) diaspore mass, B) total diaspore length, C) terminal velocity, and D) long distance dispersal potential.

|  | Estimate | Std. Error | df | t-value | p-value |
| --- | --- | --- | --- | --- | --- |
| Intercept | 0.160 | 0.086 | 103.42 | 1.861 | 0.066 |
| Habitat Continuity | -0.231 | 0.100 | 103.15 | -2.331 | 0.022 |
| DD5 | -0.011 | 0.011 | 103.35 | -1.015 | 0.313 |
| Habitat Continuity:DD5 | 0.031 | 0.013 | 103.34 | 2.347 | **0.021** |

**B) Total Diaspore Length**
|  | Estimate | Std. Error | df | t-value | p-value |
| --- | --- | --- | --- | --- | --- |
| Intercept | 13.583 | 7.505 | 100.75 | 1.810 | 0.073 |
| Habitat Continuity | -26.853 | 8.674 | 100.16 | -3.096 | 0.003 |
| DD5 | -0.927 | 0.990 | 100.75 | -0.938 | 0.351 |
| Habitat Continuity:DD5 | 3.543 | 1.000 | 100.36 | 3.093 | **0.003** |

**C) Terminal Velocity**
|  | Estimate | Std. Error | df | t-value | p-value |
| --- | --- | --- | --- | --- | --- |
| Intercept | 0.065 | 0.757 | 42.95 | 0.086 | 0.932 |
| Habitat Continuity | 2.918 | 0.983 | 41.15 | 2.968 | 0.005 |
| DD5 | 0.113 | 0.100 | 42.97 | 1.132 | 0.264 |
| Habitat Continuity:DD5 | -0.380 | 0.130 | 41.22 | -2.929 | **0.006** |

**D) Long Distance Dispersal Potential**
|  | Estimate | Std. Error | df | t-value | p-value |
| --- | --- | --- | --- | --- | --- |
| Intercept | 3.373 | 6.079 | 43.30 | 0.555 | 0.582 |
| Habitat Continuity | -18.940 | 7.917 | 41.26 | -2.392 | 0.021 |
| DD5 | -0.279 | 0.802 | 43.33 | -0.348 | 0.730 |
| Habitat Continuity:DD5 | 2.736 | 1.044 | 41.35 | 2.620 | **0.012** |

**Figure 2:**
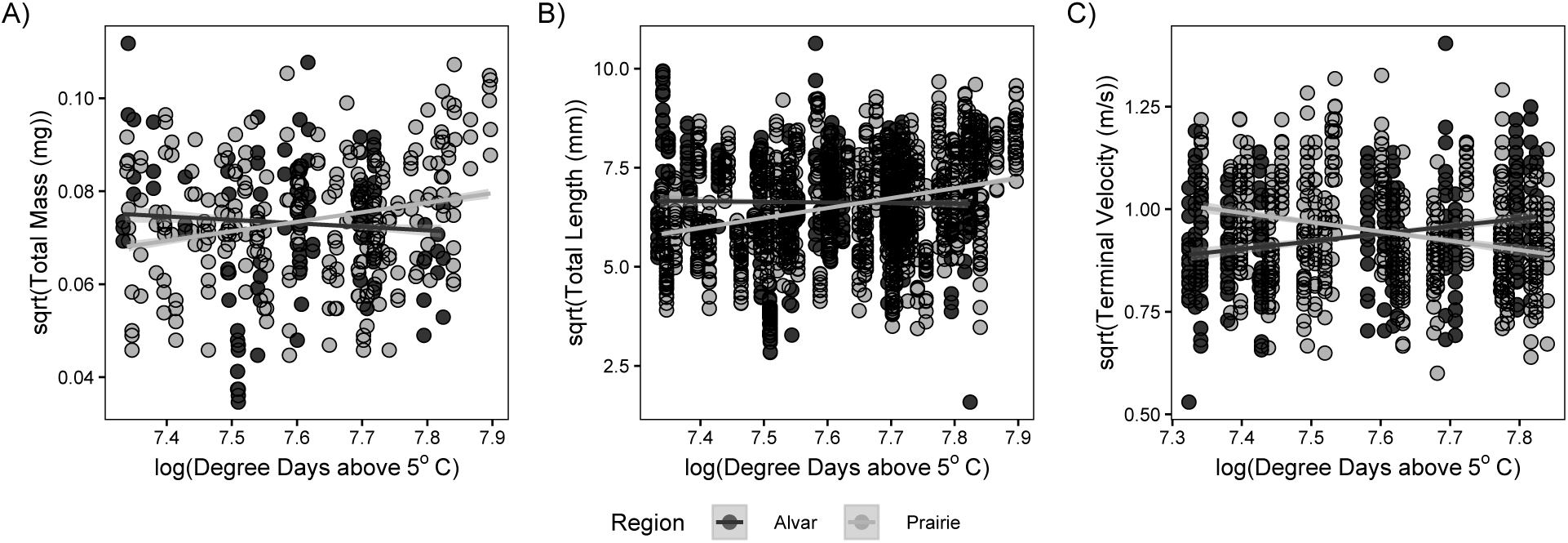
Relationship between diaspore traits and degree days above 5oC (DD5) in both prairie (light gray) and alvar (dark gray) regions. Total diaspore mass A) showed a significant interaction between habitat continuity (prairie vs alvar) and local climate – as the growing season gets longer (increased DD5), diaspore mass increased in prairies, while alvar diaspore mass decreased slightly. Diaspore length B) shows a significant interaction between habitat continuity and local climate. As the season gets longer (increased DD5), in prairies diaspores get longer, and in alvars there is little change. Terminal velocity C) showed a strong interaction between habitat continuity and local climate. As the length of the growing season increased (increased DD5), in alvars the terminal velocity of a diaspore increased, in prairies the terminal velocity decreased. This indicates that diaspores have the potential to travel further distances in prairies than in alvars in longer seasons, but travel further in alvars when the season is shorter.

### Morphology Measurements

We found a significant interaction between habitat continuity (alvar vs prairie) and local climate on total diaspore length (seed plus style length) (Table 1b, *p*=0.003), with fixed effects explaining 7.74% of the variation, and fixed and random effects together explaining 80.8% of the variation. This interaction suggests as growing season increases, diaspore length increases in prairies, but changes little in alvars (Fig. 2b). We did not find a significant effect of region or climate on either diaspore area or diaspore shape index (Appendix S2).

### Terminal Velocity Measurements

We also observed a significant interaction between habitat continuity (alvar vs prairie) and local climate for terminal velocity of the diaspores (Table 1c, *p*=0.006), with fixed effects explaining 8.9% of the variance, and fixed and random effects together explaining 55.4% of the variance. Longer growing seasons predicts reduced terminal velocities in the prairies (i.e. diaspores fall slower), and greater terminal velocities in the alvars (i.e. diaspores fall faster) (Fig. 2c).

### Estimated Dispersal Distances

We found that the interaction between habitat continuity (alvar vs prairie) and local climate significantly altered potential long-distance dispersal of *G. triflorum* (Table 1d, p=0.004), with fixed effects explaining 58.8% of the variance, and fixed and random effects together explaining 78.8% of the variance. Longer growing seasons predicts increased long-distance dispersal potential in the prairies, and reduced long-distance dispersal potential in the alvars (Fig. 3). The strength of the difference in dispersal potential between regions was likely influenced by differences in wind speed between regions, as alvars are predicted to have lower average wind speeds because of their shape (Schaefer and Larson 1997; Damschen et al. 2014). To confirm our dispersal results were robust, we re-ran the WALD model using equivalent wind conditions for both prairie and alvar individuals and still found that long-distance dispersal potential was significantly affected by the interaction between region and climate (t=3.42, *p=*0.001).

**Figure 3:**
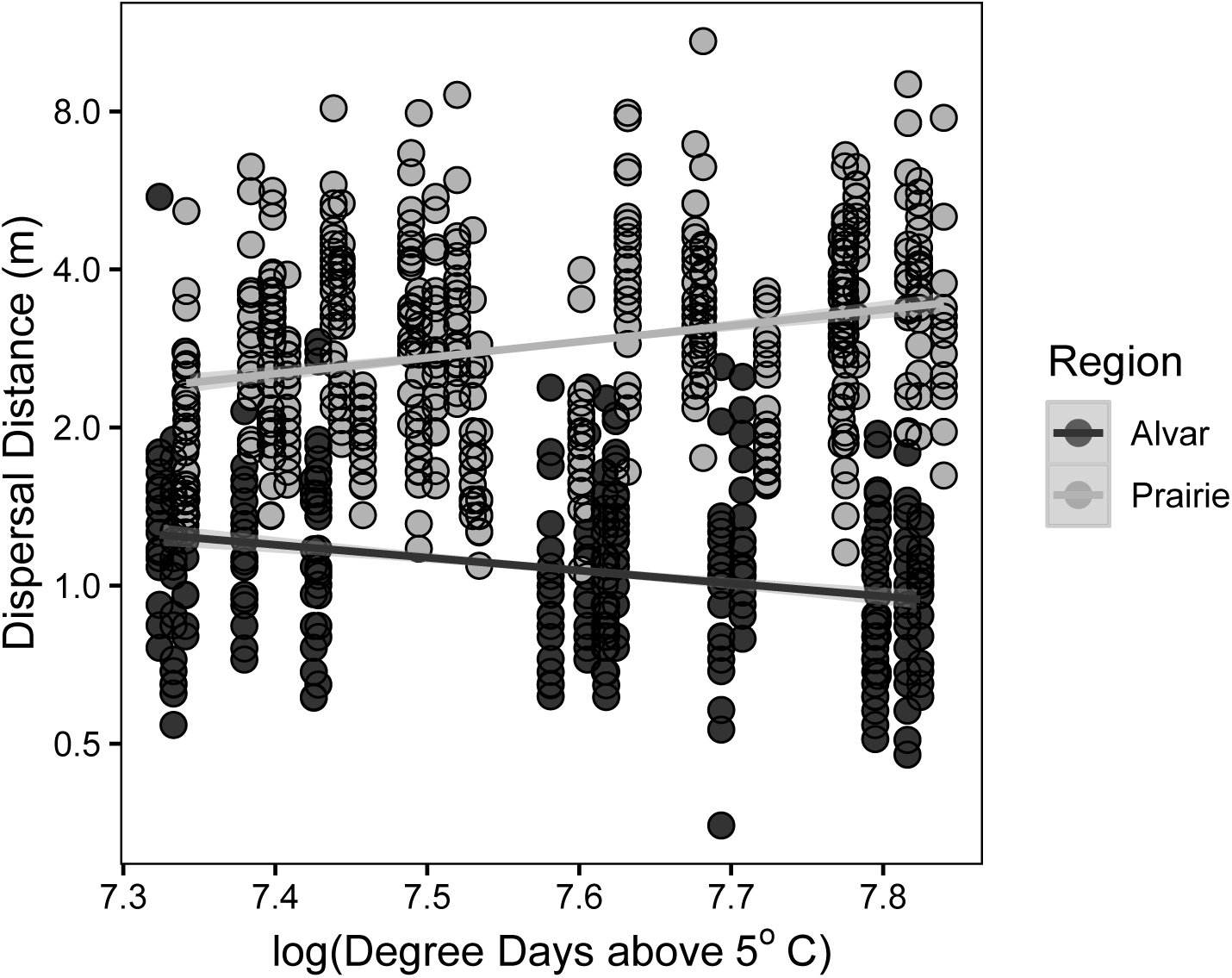
When using the WALD model to estimate dispersal distances for *G. triflorum* individuals we found a significant interaction between region and climate. As the length of the growing season increased (increased degree days above 5oC), in alvars (dark gray) the potential long-distance dispersal ability (the distance traveled by the farthest 1%) of a diaspore increased, in prairies (light gray) the long-distance dispersal ability increased.

## Discussion

Quantifying the effect local climatic conditions and habitat continuity have on dispersal trait variation has implications for predicting species’ capacity to track shifting fitness optima in rapidly changing environments (Davis and Shaw 2001; Kokko and López-Sepulcre 2006). We found, with the accumulation of growing degree days, *Geum triflorum* exhibited increased dispersal potential in continuous prairie environments. In contrast, isolated alvar populations exhibited reduced dispersal potential as the number of growing degree days increased. These findings indicate that if local environmental conditions promote a longer growing season, individuals in prairie populations may have the potential to move farther from their natal location, whereas individuals in alvar populations maximize their movement potential when conditions promote a shorter growing season. Region-specific dispersal potential may become further reinforced in a changing climate where growing seasons are predicted to change (McGinn and Shepherd 2003; Wuebbles and Hayhoe 2004; Pryor et al. 2013). These findings suggest the interaction of local climate with habitat continuity via contemporary dispersal trait variation will have substantial impacts on the maintenance of population connectivity, pointing towards the importance of considering the interaction of these factors in predicting species range shifts in response to global change.

Our results demonstrate the importance of investigating how local climate and habitat continuity interact to alter dispersal ability. Rapid anthropogenic changes are influencing both climate and habitat simultaneously (Rockström 2009; Van Den Elzen et al. 2016), yet very few studies have examined how these two global change drivers interact to alter dispersal ability (but see: Cormont et al. 2011; Mantyka-Pringle et al. 2012; Delattre et al. 2013). Delattre et al. (2013) found that the interaction between temperature and habitat structure impacted movement ability of the meadow brown butterfly (*Maniola jurtina*). In fragmented systems, movement ability was temperature-dependent. At low temperatures the butterflies exhibited reduced emigration rates, but under higher temperatures greater emigration rates. However, in continuous habitats, movement was independent of shifts in temperature. While the interaction of these factors contrasts with findings in *G. triflorum*, they suggest the relationship between movement, habitat continuity and climate are complex. We encourage continued examination of the interplay between climate and habitat continuity and its impact on dispersal potential across species, especially those of conservation concern, in order to predict shifts in gene flow and genetic connectivity across heterogeneous landscapes.

There are substantial eco-evolutionary consequences for differences observed in contemporary dispersal traits. When considering the ability for diaspores to move, terminal velocity is critical (Wilson 2000). This composite trait takes into account mass, morphology, and physical structures, such as hairs, to create differences in the length of time a diaspore can stay aloft in a column of air (Greene and Johnson 1990). Terminal velocity plays an important role in a species’ response to habitat fragmentation (Schleicher et al. 2011), and indirectly gene flow and the maintenance of connectivity across populations. Our data suggest that diaspores of *G. triflorum* will have the potential to disperse farther in prairie habitats as the number of growing degree days increase, but under similar conditions dispersal potential will be reduced in alvar habitats. In addition, we found that local climate and habitat continuity altered another important dispersal trait, achene mass. This trait plays a role in establishment once individuals have dispersed, as larger propagules tend to have a higher survival probability during establishment (Moles and Westoby 2004; Skarpaas et al. 2011). We find that achene mass and terminal velocity exhibit similar trends, as achene mass increases with increasing growing degree days in prairie habitats, but only slightly decreases with increased growing degree days in alvar habitats. When combined with terminal velocity, these results indicate that as climates warm and the number of growing degree days increases, *G. triflorum* populations in prairie habitats have the potential to disperse farther and have an increased establishment probability, while alvar populations exhibit decreased dispersal and establishment potential.

In the heterogeneous landscapes over which *G. triflorum* occurs (i.e. continuous prairies vs disjunct alvars), the spatial scale of habitat availability is highly relevant to selection on dispersal ability. For alvar versus prairie habitats the frequency and location of suitable habitats may impact dispersal trait variation. According to the theory of island biogeography (MacArthur and Wilson 1967), populations in isolated habitats may select against dispersal, particularly where long-distance dispersal leads to reduced fitness as organisms disperse beyond the range of suitable habitats (Schenk 2013; Reluga and Shaw 2015; Shaw et al. 2019). Disjunct alvar habitats resemble oceanic islands, existing as open, isolated limestone barrens situated within a matrix of boreal forest. Similar to gypsum outcrops described by Van Den Elzen et al. (2016), the low frequency of disjunct alvar habitats relative to more continuous prairies will lead to strong selection against dispersal within alvar populations, as propagules that disperse longer distances likely land in unsuitable habitat. While selection may have shifted to favor reduced dispersal for isolated alvar populations, the same may not be true of prairie populations. Relatively continuous prairie habitats increases the likelihood that dispersing propagules will land in suitable habitat, contributing to selection for increased dispersal (Travis and Dytham 1999). However, previous and ongoing agricultural conversion may increase the scale of isolation for populations within a matrix of inhospitable environments (Wright and Wimberly 2013; Lark et al. 2018; Wimberly et al. 2018). Thus, increasing fragmentation across prairie environments has the potential to shift the direction of selection for dispersal traits in these historically continuous habitats.

The colonization history of isolated alvar habitats suggests that the strength and direction of selection for dispersal traits for grassland species, such as *G. triflorum*, isolated on alvar habitats could have changed over time (Catling and Brownell 1995; Hamilton and Eckert 2007). Previous research suggests that alvar habitats were likely colonized by grassland species during the warming Hypsithermal period ~5000 YBP as grasslands expanded (Hamilton and Eckert 2007). However, following a cooling period, subsequent range contractions likely isolated a number of grassland species, including *G. triflorum*, on alvar habitats (Catling and Brownell 1995; Hamilton and Eckert 2007). While traits associated with long-distance dispersal may have been selected for during initial colonization of alvar habitats, following isolation selection may act against traits associated with dispersal, as our results suggest. Furthermore, ecological specialization and local adaptation across alvar habitats may limit further gene flow, strengthening selection against dispersal (Yoko et al. n.d.; Van Den Elzen et al. 2016). In order to determine the potential strength of selection in these two different habitats, common garden experiments may be used to tease apart the respective role of the environmental variation from genetic differences, which reflect both heritable genetic differences and maternal effects, both of which have been shown to alter dispersal traits (Yoko et al. n.d.; Donohue 1998; Galloway 2005; Jacobs and Lesmeister 2012). In addition, while we have captured a snapshot of the relationship between environment and dispersal trait variation, temporal monitoring of trait variation within reciprocal transplant experiments will be essential to understanding the role plasticity may play in maintaining variation in dispersal traits in response to seasonal environmental change.

Local climatic variation and shifting habitat suitability have the potential to rapidly alter dispersal potential, impacting predictions for connectivity and species’ range shifts in response to global change (Kubisch et al. 2013). Where suitable habitat is continuous, increased dispersal potential may facilitate range expansion and connectivity under favorable climates, however, the impact of spatial isolation could increase in a warming climate, exacerbating the consequences of isolation. This may impact the evolutionary potential of populations, particularly those isolated populations at the periphery of a species’ range where reduced connectivity influences the maintenance of genetic variation (Jump and Peñuelas 2005). Thus, species’ ability to sustain an evolutionary response to changing conditions requires an evaluation of the interaction between climate and landscape structure to establish predictions for species distributions and connectivity under global change.

## Acknowledgements

We thank Chris Eckert for support during early collections, and Jon Sweetman, Tyler Stadel, Steve Travers, and Rebekah Neufeld for help field sampling, Grace Tharmarajah, Pat Welsh, Stephen Johnson, and Jeffrey Kittilson for lab help and diaspore measurements, and Adam Clark for consultation on the terminal velocity measurer. We particularly thank Frank Baker, Melody Kroll, Bruce McClure, Kate Wynne and the Molecular Cytology Core at the University of Missouri for help with the diaspore images, as well as Bruce Ford with the University of Manitoba (WIN) herbaria, the National Collection of Vascular Plants (DAO), and Canadian Museum of Nature (CAN) for providing high-resolution scans of herbarium specimens for use in modeling dispersal. We also thank Chris Catano, Amanda Koltz, Jonathan Myers, and Dilys Vela for productive feedback on the manuscript. L.L.S was supported by the LCCMR ENRTF Grant (M.L. 2016, Chp. 186, Sec. 2, Subd. 08b), and startup provided by the University of Missouri. J.A.H. was supported by a New Faculty award from the office of the North Dakota Experimental Program to Stimulate Competitive Research (ND-EPSCoR NSF-IIA-1355466) and startup provided by North Dakota State University.

## References

Aitken, S. N., S. Yeaman, J. A. Holliday, T. Wang, and S. Curtis-McLane. 2008. Adaptation, migration or extirpation: Climate change outcomes for tree populations. Evolutionary Applications 1:95–111.

Baguette, M., D. Legrand, J. Freville, J. Van Dyck, and S. Ducatez. 2012. Evolutionary ecology of dispersal in fragmented landscape. Pages 381–391 inDispersal ecology and evolution. Oxford University Press.

Barton, K. 2018. MuMIn: Multi-Model Inference. R package version 1.40.4.

Bates, D., M. Machler, B. Bolker, and S. Walker. 2015. Fitting linear mixed-effects models using lme4. Journal of Statistical Software 67:1–48.

Beaubien, E., and A. Hamann. 2011. Spring flowering response to climate change between 1936 and 2006 in Alberta, Canada. BioScience 61:514–524.

Bjorkman, A. D., I. H. Myers-Smith, S. C. Elmendorf, S. Normand, N. Rüger, P. S. A. Beck, A. Blach-Overgaard, et al. 2018. Plant functional trait change across a warming tundra biome. Nature 562:57–62.

Brehm, A. M., A. Mortelliti, G. A. Maynard, and J. Zydlewski. 2019. Land use change and the ecological consequences of personality in small mammals. Ecology Letters (IN PRESS).

Catling, P. M., and V. R. Brownell. 1995. A review of the alvars of the Great Lakes region: Distribution, floristic composition, biogeography and protection. Canadian Field-Naturalist 109:143–171.

Cheptou, P.-O., O. Carrue, S. Rouifed, and A. Cantarel. 2008. Rapid evolution of seed dispersal in an urban environment in the weed Crepis sancta. Proceedings of the National Academy of Sciences of the United States of America 105:3796–3799.

Cody, M. L., and J. M. Overton. 1996. Short-term evolution of reduced dispersal in island plant populations. Journal of Ecology 84:53–61.

Corlett, R. T., and D. a Westcott. 2013. Will plant movements keep up with climate change? Trends in ecology & evolution 28:482–8.

Cormont, A., A. H. Malinowska, O. Kostenko, V. Radchuk, L. Hemerik, M. F. WallisDeVries, and J. Verboom. 2011. Effect of local weather on butterfly flight behaviour, movement, and colonization: Significance for dispersal under climate change. Biodiversity and Conservation 20:483–503.

Damschen, E. I., D. V Baker, G. Bohrer, R. Nathan, J. L. Orrock, J. R. Turner, L. A. Brudvig, et al. 2014. How fragmentation and corridors affect wind dynamics and seed dispersal in open habitats. Proceedings of the National Academy of Science 111:3484–9.

Davis, M. B., and R. G. Shaw. 2001. Range shifts and adaptive responses to Quaternary climate change. Science (New York, N.Y.) 292:673–9.

Delattre, T., M. Baguette, F. Burel, V. M. Stevens, H. Quénol, and P. Vernon. 2013. Interactive effects of landscape and weather on dispersal. Oikos 122:1576–1585.

Donohue, K. 1998. Maternal determinants of seed dispersal in Cakile edentula: Fruit, plant, and site traits. Ecology 79:2771–2788.

Edelaar, P., and D. I. Bolnick. 2012. Non-random gene flow: An underappreciated force in evolution and ecology. Trends in Ecology and Evolution 27:659–665.

Eriksson, O., and A. Jakobsson. 1999. Recruitment trade-offs and the evolution of dispersal mechanisms in plants. Evolutionary Ecology 13:411–423.

Fahrig, L. 2003. Effects of Habitat Fragmentation on Biodiversity. Annual Review of Ecology, Evolution, and Systematics 34:487–515.

Fahrig, L. 2017. Ecological responses to habitat fragmentation perse. Annual Review of Ecology, Evolution, and Systematics 48:1–23.

Gajewski, W. 1958. Evolution in the genus Geum. Evolution 13:378–388.

Galloway, L. F. 2005. Maternal effects provide phenotypic adaptation to local environmental conditions. The New Phytologist 166:93–9.

Greene, D. F., and E. A. Johnson. 1989. A model of wind dispersal of winged or plumed seeds. Ecology 70:339–347.

Greene, D. F., and E. A. Johnson. 1990. The aerodynamics of plumed seeds. Functional Ecology 4:117–125.

Hamilton, J. A., and C. G. Eckert. 2007. Population genetic consequences of geographic disjunction: A prairie plant isolated on Great Lakes alvars. Molecular Ecology 16:1649–1660.

Hamilton, W. D., and R. M. May. 1977. Dispersal in stable habitats. Nature 269:578–581.

Hargreaves, A. L., S. F. Bailey, and R. A. Laird. 2015. Fitness declines towards range limits and local adaptation to climate affect dispersal evolution during climate-induced range shifts. Journal of Evolutionary Biology 28:1489–1501.

Hargreaves, A. L., and C. G. Eckert. 2014. Evolution of dispersal and mating systems along geographic gradients: Implications for shifting ranges. Functional Ecology 28:5–21.

Higgins, S. I., O. Flores, and F. M. Schurr. 2008. Costs of persistence and the spread of competing seeders and sprouters. Journal of Ecology 96:679–686.

Jacobs, B. S., and S. A. Lesmeister. 2012. Maternal environmental effects on fitness, fruit morphology and ballistic seed dispersal distance in an annual forb. Functional Ecology 26:588–597.

Jump, A. S., and J. Peñuelas. 2005. Running to stand still: Adaptation and the response of plants to rapid climate change. Ecology Letters 8:1010–1020.

Katul, G. G., A. Porporato, R. Nathan, M. Siqueira, M. B. Soons, D. Poggi, H. S. Horn, et al. 2005. Mechanistic analytical models for long-distance seed dispersal by wind. The American Naturalist 166:368–381.

Kokko, H., and A. López-Sepulcre. 2006. From individual dispersal to species ranges: Perspectives for a changing world. Science (New York, N.Y.) 313:789–791.

Kubisch, A., T. Degen, T. Hovestadt, and H. J. Poethke. 2013. Predicting range shifts under global change: The balance between local adaptation and dispersal. Ecography 36:873–882.

Kuznetsova, A., P. Bruun Brockhoff, and R. Haubo Bojesen Christensen. 2014. lmerTest: Tests for random and fixed effects for linear mixed effect models (lmer objects of lme4 package).

Lark, T. J., B. Larson, I. Schelly, S. Batish, and H. K. Gibbs. 2018. Accelerated conversion of native prairie to cropland in Minnesota. Environmental Conservation 1–8.

Lentink, D., W. B. Dickson, J. L. van Leeuwen, and M. H. Dickinson. 2009. Leading-edge vortices elevate lift of autorotating plant seeds. Science (New York, N.Y.) 324:1438–40.

MacArthur, R. H., and E. O. Wilson. 1967. The theory of island biogeography. Princeton University Press, New Jersey.

Mantyka-Pringle, C. S., T. G. Martin, and J. R. Rhodes. 2012. Interactions between climate and habitat loss effects on biodiversity: A systematic review and meta-analysis. Global Change Biology 18:1239–1252.

McGinn, S. M., and A. Shepherd. 2003. Impact of climate change scenarios on the agroclimate of the Canadian prairies. Canadian Journal of Soil Science 83:623–630.

Moles, A. T., and M. Westoby. 2004. Seedling survival and seed size: a synthesis of the literature. Journal of Ecology 92:372–383.

Parmesan, C. 2006. Ecological and evolutionary responses to recent climate change. Annual Review of Ecology, Evolution, and Systematics 37:637–669.

Parmesan, C., and G. Yohe. 2003. A globally coherent fingerprint of climate change impacts across natural systems. Nature 421:37–42.

Pryor, S. C., R. J. Barthelmie, and J. T. Schoof. 2013. High-resolution projections of climate-related risks for the Midwestern USA. Climate Research 56:61–79.

R Core Team. 2018. R: A Language and Environment for Statistical Computing.

Reluga, T. C., and A. K. Shaw. 2015. Resource distribution drives the adoption of migratory, partially migratory, or residential strategies. Theoretical Ecology 8:437–447.

Reschke, C., R. Reid, J. Jones, T. Feeny, and H. Potter. 1999. Conserving Great Lakes Alvars. Final technical report of the international alvar conservation initiative. Chicago.

Rockström, J. 2009. A safe operating space for humanity. Nature 461:472–475.

Rohrer, J. R. 1993. Geum. Pages 1–10. in F. of N. A. E. Committee, ed. Flora of North America North of Mexico. 20+ vols. (Vol. 9). New York and Oxford.

Schaefer, C. A., and D. W. Larson. 1997. Vegetation, environmental characteristics and ideas on the maintenance of alvars on the Bruce Peninsula, Canada. Journal of Vegetation Science 8:797–810.

Schenk, J. J. 2013. Evolution of limited seed dispersal ability on gypsum islands. American Journal of Botany 100:1811–22.

Schleicher, A., R. Biedermann, and M. Kleyer. 2011. Dispersal traits determine plant response to habitat connectivity in an urban landscape. Landscape Ecology 26:529–540.

Schneider, C. A., W. S. Rasband, and K. W. Eliceiri. 2012. NIH Image to ImageJ: 25 years of image analysis. Nature Methods 9:671.

Sexton, J. P., P. J. McIntyre, A. L. Angert, and K. J. Rice. 2009. Evolution and ecology of species range limits. Annual Review of Ecology, Evolution, and Systematics 40:415–436.

Shaw, A. K., C. C. D’Aloia, and P. M. Buston. 2019. The evolution of marine larval dispersal kernels in spatially structured habitats: Analytical models, individual-based simulations, and comparisons with empirical estimates. The American Naturalist 3:424–435.

Skarpaas, O., E. J. SIlverman, E. Jongejans, and K. Shea. 2011. Are the best dispersers the best colonizers? Seed mass, dispersal and establishment in Carduus thistles. Evolutionary Ecology 25:155–169.

Soons, M. B., G. W. Heil, R. Nathan, and G. G. Katul. 2004. Determinants of long-distance seed dispersal by wind in grasslands. Ecology 85:3056–3068.

Sullivan, L. L., A. T. Clark, D. Tilman, and A. K. Shaw. 2018. Mechanistically derived dispersal kernels explain species-level patterns of recruitment and succession. Ecology 99:2415–2420.

Teller, B. J., C. Campbell, and K. Shea. 2014. Dispersal under duress: Can stress enhance the performance of a passively dispersed species? Ecology 95:2694–2698.

Thomson, F. J., A. T. Moles, T. D. Auld, and R. T. Kingsford. 2011. Seed dispersal distance is more strongly correlated with plant height than with seed mass. Journal of Ecology 99:1299–1307.

Tilman, D. 1994. Competition and biodiversity in spatially structured habitats. Ecology 75:2–16.

Travis, J. M. J., and C. Dytham. 1998. The evolution of dispersal in a metapopulation: a spatially explicit, individual-based model. Proceedings of the Royal Society B: Biological Sciences 265:17–23.

Travis, J. M. J., and C. Dytham. 1999. Habitat persistence, habitat availability and the evolution of dispersal. Proceedings of the Royal Society B: Biological Sciences 266:723–728.

Van Den Elzen, C. L., E. A. LaRue, and N. C. Emery. 2016. Oh, the places you’ll go! Understanding the evolutionary interplay between dispersal and habitat adaptation as a driver of plant distributions. American Journal of Botany 103:2015–2018.

Varshney, K., S. Chang, and Z. J. Wang. 2012. The kinematics of falling maple seeds and the initial transition to a helical motion. Nonlinearity 25:C1–C8.

Wang, T., A. Hamann, D. Spittlehouse, and C. Carroll. 2016. Locally downscaled and spatially customizable climate data for historical and future periods for North America. PLoS ONE 11:1–17.

Whittet, R., S. Cavers, J. Cottrell, C. Rosique-Esplugas, and R. Ennos. 2017. Substantial variation in the timing of pollen production reduces reproductive synchrony between distant populations of Pinus sylvestris L. in Scotland. Ecology and Evolution 7:5754–5765.

Williams, J. L., B. E. Kendall, and J. M. Levine. 2016. Rapid evolution accelerates plant population spread in fragmented experimental landscapes. Science (New York, N.Y.) 353:482–485.

Wilson, J. D. 2000. Trajectory models for heavy particles in atmospheric turbulence: Comparison with observations. Journal of Applied Meteorology 39:1894–1912.

Wimberly, M. C., D. M. Narem, P. J. Bauman, B. T. Carlson, and M. A. Ahlering. 2018. Grassland connectivity in fragmented agricultural landscapes of the north-central United States. Biological Conservation 217:121–130.

Wright, C. K., and M. C. Wimberly. 2013. Recent land use change in the Western Corn Belt threatens grasslands and wetlands. Proceedings of the National Academy of Science 110:4134–4139.

Wuebbles, D. J., and K. Hayhoe. 2004. Climate change projections for the United States Midwest. Mitigation and Adaptation Strategies for Global Change 9:335–363.

Yoko, Z., K. Volk, N. A. Dochtermann, and J. A. Hamilton. n.d. Evolution of quantitative trait differentiation across heterogeneous landscapes.

Zhu, K., C. W. Woodall, and J. S. Clark. 2012. Failure to migrate: Lack of tree range expansion in response to climate change. Global Change Biology 18:1042–1052.

Zobel, M., M. Moora, and T. Herben. 2010. Clonal mobility and its implications for spatio-temporal patterns of plant communities: What do we need to know next? Oikos 119:802–806.

